# Increased Netrin downstream of overactive Hedgehog signaling disrupts optic fissure formation

**DOI:** 10.1101/2024.06.18.599642

**Authors:** Sarah Lusk, Sarah LaPotin, Jason S. Presnell, Kristen M. Kwan

**Affiliations:** Department of Human Genetics, University of Utah, Salt Lake City, UT 84112

**Keywords:** *netrin1a*, coloboma, eye development, morphogenesis, *ptch2*

## Abstract

**Background:** Uveal coloboma, a developmental eye defect, is caused by failed development of the optic fissure, a ventral structure in the optic stalk and cup where axons exit the eye and vasculature enters. The Hedgehog (Hh) signaling pathway regulates optic fissure development: loss-of-function mutations in the Hh receptor *ptch2* produce overactive Hh signaling and can result in coloboma. We previously proposed a model where overactive Hh signaling disrupts optic fissure formation by upregulating transcriptional targets acting both cell- and non-cell-autonomously. Here, we examine the Netrin family of secreted ligands as candidate Hh target genes.

**Results:** We find multiple Netrin ligands upregulated in the zebrafish *ptch2* mutant during optic fissure development. Using a gain-of-function approach to overexpress Netrin in a spatiotemporally specific manner, we find that *netrin1a* or *netrin1b* overexpression is sufficient to cause coloboma and disrupt wild-type optic fissure formation. We used loss-of-function alleles, CRISPR/Cas9 mutagenesis, and morpholino knockdown to test if loss of Netrin can rescue coloboma in the *ptch2* mutant: loss of *netrin* genes does not rescue the *ptch2* mutant phenotype.

**Conclusion:** These results suggest that Netrin is sufficient but not required to disrupt optic fissure formation downstream of overactive Hh signaling in the *ptch2* mutant.

**Key Findings:**

- Overactive Hedgehog signaling in the *ptch2* mutant causes increased netrin expression
- Spatiotemporally specific overexpression of netrin1a and netrin1b can cause coloboma
- Spatiotemporally specific overexpression of netrin1a can disrupt optic fissure and stalk formation as well as optic stalk cell morphology, similar to the *ptch2* mutant
- Loss of netrin ligands in the *ptch2* mutant does not rescue the phenotype

## Introduction

The proper three-dimensional structure of the eye is critical for vision, as structural defects commonly account for visual impairment in newborns. One such defect, uveal coloboma, is caused by failed development of the optic fissure, a transient seam along the ventral surface of the optic stalk and optic cup that forms a conduit during development for retinal ganglion cell axons to exit the eye and vasculature to enter. Uveal coloboma is a significant cause of pediatric blindness worldwide, yet we lack a basic understanding of the cellular and molecular mechanisms disrupted ^1–5^. Through human genetic studies and findings using animal models, we know that the genetic underpinnings of coloboma are heterogeneous and include mutations in multiple signaling pathways ^4,6^.

One pathway central to optic fissure development is the Hedgehog (Hh) signaling pathway: mutations upstream, within, and downstream of this pathway can result in coloboma ^4^. Human mutations in the Hh receptor *PTCH* cause Gorlin syndrome ^7,8^, in which affected individuals are typically diagnosed with medulloblastoma or basal cell carcinoma, along with numerous additional phenotypes including coloboma ^9^.

In zebrafish, the *ptch2* loss-of-function mutant displays coloboma: given the function of Ptch2 as a negative regulator of Hh signaling, these mutations lead to overactive Hh signaling. The *ptch2* mutant coloboma phenotype has been described in detail ^10^, however, molecular mechanisms directly driving disruption of optic fissure morphogenesis are still unclear. We previously determined the cellular mechanisms by which the *ptch2* mutant phenotype initially arises ^11^. Using multidimensional timelapse microscopy and cell tracking, we identified the cells that give rise to the optic fissure in wild-type embryos. In the *ptch2* mutant, these cells do not move to their correct position; as a result, the optic fissure fails to form. Additional analyses of cells in the optic stalk revealed morphological defects at the single cell level: cells are less elongated compared to the wild-type optic stalk. Downstream transcriptional targets are upregulated in the *ptch2* mutant, and Gli activity is required for the mutant phenotype. In addition, the *ptch2* mutant phenotype is regulated by both cell autonomous and non-cell autonomous mechanisms ^11^. Taken together, these data suggest that overactive Hh signaling in the *ptch2* mutant disrupts cell movements and morphology via misregulation of downstream transcriptional targets acting intra- and inter-cellularly, resulting in aberrant optic fissure and stalk formation. We hypothesize that this disruption of optic fissure and stalk formation underlies the *ptch2* mutant coloboma phenotype.

Therefore, downstream transcriptional targets of Hh signaling are likely the key factors that directly disrupt optic fissure and stalk cell movements and cause coloboma. To identify these downstream factors, we have taken a candidate approach, focusing initially on intercellular signaling molecules that are known transcriptional targets of Hh signaling and are expressed at the appropriate time and place to influence optic fissure and stalk morphogenesis. Using these criteria, we have identified an initial candidate: Netrin, a family of laminin-related secreted molecules, largely studied in the context of axon guidance. In this study we examine the genetic interaction between *netrin* and the Hh signaling pathway to determine if upregulation of Netrin is, in part, responsible for the *ptch2* mutant coloboma phenotype.

Netrin family proteins are diffusible molecules that can regulate diverse developmental processes, including but not limited to axonal guidance, cell survival, and cell-cell adhesion ^12–15^. Zebrafish contain five *netrin* genes: *netrin 1a* (*ntn1a*), *netrin 1b* (*ntn1b*), *netrin 2* (*ntn2*), *netrin 4* (*ntn4*), and *netrin 5* (*ntn5*) ^16–19^. Roles for Netrin1 in eye development have previously been described in zebrafish, chick and mouse. For example, Ntn1a acts as a retinal ganglion cell axon guidance molecule expressed along the optic fissure ^20^. However, *ntn1a* is also expressed at an earlier stage in the nasal optic vesicle and optic vesicle junction with the forebrain ^21^. This suggests that *ntn1a* is expressed at the right time and in the right location to act downstream of Hh signaling in optic fissure and stalk formation. Further, both *ntn1a* and *ntn1b* expression have been established as being responsive to Hh signaling: in *sonic hedgehog a* (*shha*) and *smoothened* (*smo*) mutants, both of which have decreased Hh signaling, *ntn1a* and *ntn1b* mRNA levels are decreased ^22,23^. In response to increased Hh signaling, for example, *sonic Hh* (*Shh*) and a dominant negative form of *protein kinase A* (*dnPKA*) overexpression, *ntn1a* expression is ectopically expanded in all regions, including the head and eyes ^24,25^. Although *ntn1a* expression has been assayed at timepoints relevant to optic fissure and stalk formation, other Netrins have only been analyzed at later stages ^16,18,26^. Of interest, two recent studies utilized optic fissure transcriptomic approaches to identify novel coloboma causing genes and found Netrin1 mediates optic fissure closure later in development ^27,28^.

Here, we characterize a novel role for Netrin during early eye development: we asked whether *netrin* might be a key downstream target of Hedgehog signaling in the *ptch2* mutant, in which overactive signaling leads to defective optic fissure formation and coloboma. We find that in wild-type embryos, upregulation of *netrin* in Hh-responding cells is sufficient to disrupt optic fissure formation and can lead to coloboma. Despite finding that spatiotemporally specific overexpression of *netrin* is sufficient to lead to coloboma, these ligands are not required for the *ptch2* mutant phenotype; removal of multiple *netrin* ligands from the *ptch2* mutant is unable to rescue coloboma. This suggests that although Netrin may be involved in both formation and fusion of the optic fissure, there is likely functional redundancy in this complex morphological defect, particularly in the formation step. Together, this work uncovers molecular mechanisms regulating optic fissure formation and, in turn, coloboma.

## Results

### netrin genes are expressed in the head during optic cup morphogenesis and are upregulated in response to overactive Hh signaling

Research in different animal models has demonstrated that *netrin* genes are expressed widely during development and in a variety of organs ^12,14,29–34^. In zebrafish, it has been previously shown that both *ntn1a* and *ntn1b* are expressed in the optic vesicle ^21,22^, however it remains unknown if *ntn2*, *ntn4*, or *ntn5* are expressed during early eye development in zebrafish. Thus, we first examined which ligands are expressed in the early embryo to determine which might regulate optic fissure morphogenesis. We performed whole mount *in situ* hybridization for all 5 genes in 24 hours post fertilization (hpf) wild-type embryos (**Fig. 1A-E’**). We found that *ntn1a* and *ntn1b* are expressed in a distinct pattern restricted to the embryonic midline, and potentially within the optic stalk (**Fig. 1A-B’**). *ntn2*, *ntn4*, and *ntn5* appear lowly expressed and not spatially restricted at this stage (**Fig. 1C-E’**).

**Figure 1.**
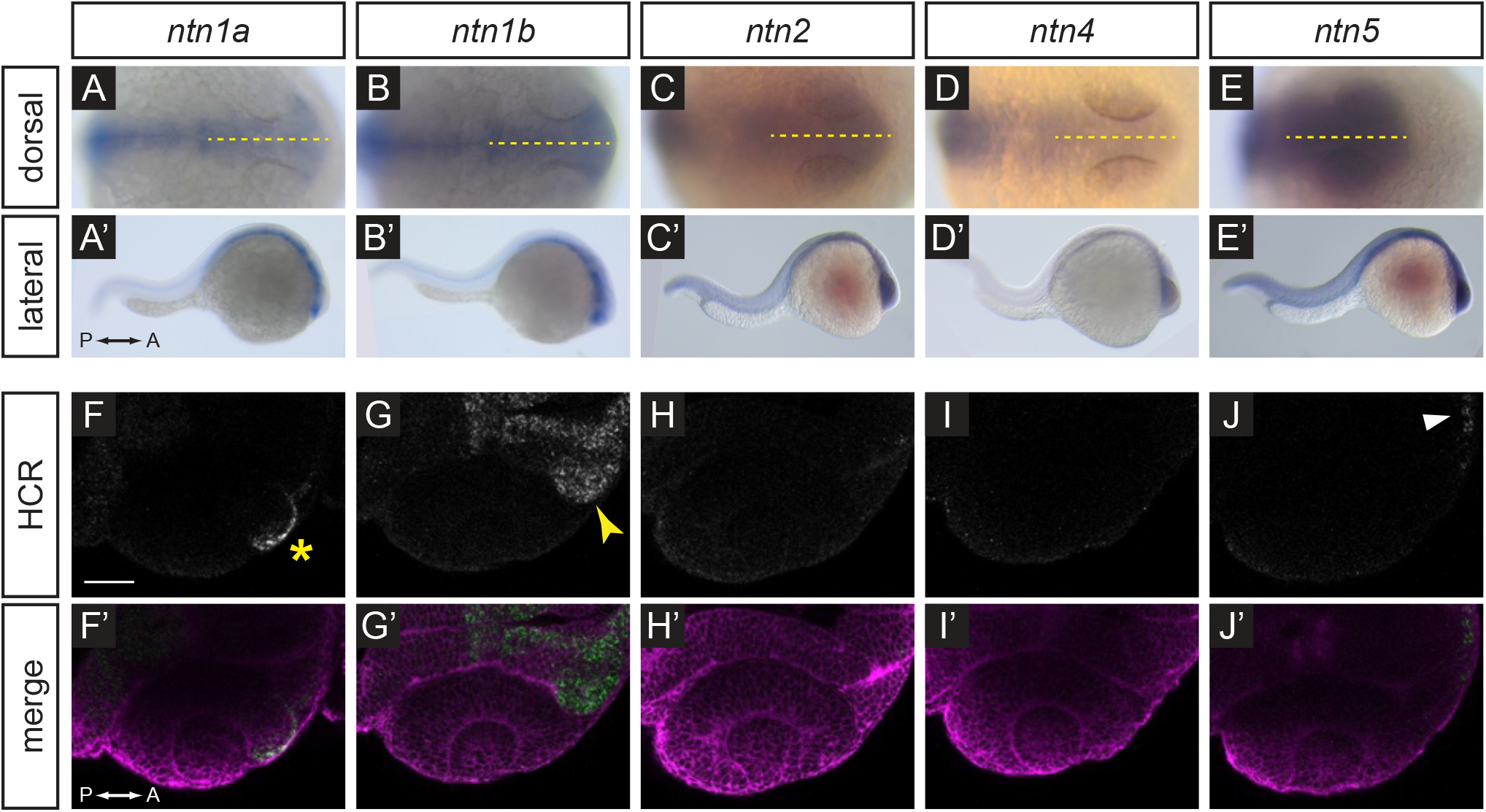
Expression patterns of *netrin* ligands. (A-E’) Whole-mount *in situ* hybridization for *ntn1a, ntn1b, ntn2, ntn4, and ntn5* in wild-type embryos at 24 hpf, (A-E) dorsal orientation, (A’-E’) lateral orientation. Yellow dotted lines (A-E) indicate embryo midline. (F-J’) HCR RNA-FISH for *ntn1a, ntn1b, ntn2, ntn4, and ntn5* in wild-type embryos harboring the *Tg(bactin2:EGFP-CAAX)* transgene at 24 hpf, (F-J) HCR signal alone. (F’-J’) Merge of HCR (green) and *Tg(bactin2:EGFP-CAAX)* transgene (magenta) to visualize tissue morphology. Yellow asterisk (F) indicates *ntn1a* signal in the nasal margin of the optic fissure; yellow arrowhead (G) indicates *ntn1b* signal in the optic stalk; white arrowhead (J) indicates faint *ntn5* in the telencephalon. Scale bar, 50 µm.

In order to visualize gene expression with greater spatial resolution, we utilized hybridization chain reaction RNA fluorescence *in situ* (HCR RNA-FISH) technology ^35^. Probe target sites were selected to be specific to each *netrin* gene, and HCR was performed on embryos fixed at 24 hpf from a *ptch2^+/-^*; *Tg(bactin2:EGFP-CAAX)* incross. In wild-type embryos, we observe a similar expression pattern for *ntn1a* and *ntn1b* at this stage as we did with colorimetric *in situ* hybridization. *ntn1a* is expressed in the midline, in addition to the anterior rim of the optic cup and nasal margin of the optic fissure (**Fig. 1F-F’**; yellow asterisk marks nasal margin of the fissure). *ntn1b* is expressed in a broader pattern than *ntn1a* and includes expression within the optic stalk, but not the optic cup (**Fig. 1G-G’**; yellow arrowhead indicates stalk). Expression of *ntn2* and *ntn4* is not detected within the head at this stage (**Fig. 1H-I’**). *ntn5* expression appears restricted to only the few most anterior, ventral cells within the forebrain (**Fig. 1J-J’**; white arrowhead indicates forebrain).

To determine if Netrin ligands are responsive to Hh signaling we assayed gene expression in *ptch2* mutants and siblings. We observe that *ntn1a* expression is apparent in the optic fissure in *ptch2^-/-^* embryos similar to wild-type, however expression appears expanded (**Fig. 2A-A’’’, B-B’’’**; asterisks indicate expression in the nasal optic fissure). We quantified expression by measuring the domain angle in degrees (schematized in **Fig. 2E**) and found that indeed the domain of expression is expanded in *ptch2^-/-^* compared to wild-type siblings (**Fig. 2E**; sibling 78.69 ± 7.51°; *ptch2^-/-^* 124.5 ±15.08°). Similar to wild-type, *ntn1b* expression in *ptch2^-/-^* is observed in the optic stalk, but expression appears increased (**Fig. 2C-C’’’, D-D’’’**; arrowheads indicate stalk). We further measured *ntn1a* and *ntn1b* expression in wild-type and *ptch2*^-/-^ embryos by quantifying and normalizing fluorescence intensity in the nasal margin of the optic fissure (*ntn1a)*, or the optic stalk (*ntn1b*); see Methods for details. We observe a statistically significant increase in *ntn1a* and *ntn1b* expression in *ptch2*^-/-^ embryos compared to wild-type siblings (**Fig. 2F**). *ntn2* and *ntn4* expression remain absent in the *ptch2*^-/-^ head at 24 hpf, unchanged from wild-type (**data not shown**). *ntn5* expression appears stronger and slightly expanded in *ptch2*^-/-^, although this is still restricted to a narrow region in the forebrain, distant from the optic fissure and stalk (**data not shown**). These data suggest that Ntn2, Ntn4, and Ntn5 may be less likely to regulate optic fissure formation.

**Figure 2.**
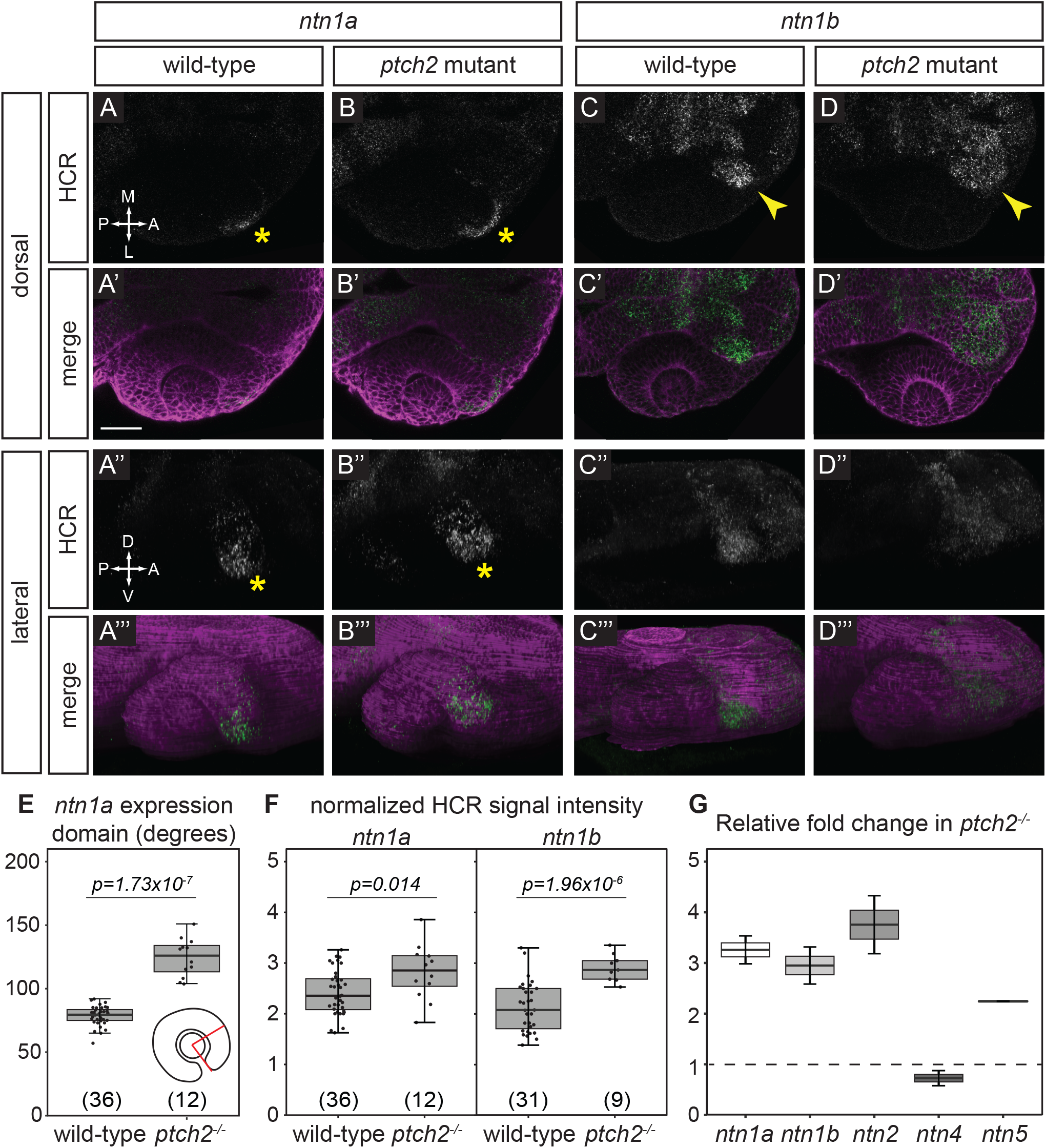
Responsiveness of *netrin* ligand expression to Hedgehog signaling. (A-D’’’) HCR RNA-FISH for *ntn1a* and *ntn1b* in wild-type embryos (A-A’’’, C-C’’’) and *ptch2^-/-^* embryos (B-B’’’, D-D’’’) at 24 hpf. Dorsal view (A-D, A’-D’) and lateral views of 3D renderings (A’’-D’’, A’’’-D’’’). The merged images (A’-D’, A’’’-D’’’) are HCR signal (green) and *Tg(bactin2:EGFP-CAAX)* transgene (magenta) to visualize tissue morphology. Yellow asterisks (A, A’’, B, B’’) indicate *ntn1a* signal in the nasal margin of the optic fissure, and yellow arrowheads (C, D) indicate *ntn1b* signal in the optic stalk. (E, F) Quantification of HCR data, for (E) the extent of the *ntn1a* expression domain; and (F) normalized fluorescence intensity for *ntn1a* in the optic fissure and *ntn1b* in the optic stalk. (G) RT-qPCR quantification showing relative fold change of *netrin* genes in *ptch2^-/-^* embryos compared to wild-type embryos at 24 hpf using the ΔΔCt method, where the relative quantity of each gene was normalized to the reference gene *eef1a1l1*. Scale bar, 50 µm.

As further evidence for increased *netrin* gene expression in *ptch2*^-/-^ embryos, we performed reverse transcription-quantitative PCR (RT-qPCR) in whole embryos at 24 hpf. We calculated the fold change in gene expression for each netrin gene normalized to a conventional housekeeping gene, *eef1a1l1,* in *ptch2*^-/-^ embryos compared to wild-type using the ΔΔCt method ^36^. We find that expression of *ntn1a*, *ntn1b*, *ntn2*, and *ntn5* genes are each at least 2-fold upregulated in *ptch2^-/-^*embryos at 24 hpf (**Fig. 2G**). Together with our findings from *in situ* hybridization, we conclude that *ntn1a* and *ntn1b* are expressed in the optic fissure and stalk respectively and are upregulated in response to overactive Hh signaling. Therefore, moving forward, we examined potential roles for *ntn1a* and *ntn1b* in optic fissure formation.

### Overexpression of Netrin in a spatiotemporally specific manner is sufficient to disrupt optic fissure formation and cause coloboma

We next asked whether Netrin overexpression might be sufficient to disrupt optic fissure formation and lead to coloboma, thereby phenocopying the *ptch2* mutant. This would suggest that Netrin might be a key downstream target of overactive Hh signaling to cause coloboma. In order to test whether overexpression of Netrin is sufficient to disrupt optic fissure and stalk formation and cause coloboma, we used the Tol2kit system ^37^, which permits transient transgenesis in injected embryos via *Tol2* transposon-mediated insertion. We used the Hh-responsive promoter *GBS-ptch2* ^38^ to overexpress Ntn1a or Ntn1b. The GBS-*ptch2* promoter restricts overexpression to cells responding to Hh ligand, the cell population that is defective in the *ptch2* mutant ^11^. This construct additionally contains a viral 2A peptide followed by a fluorescent marker, nuclear-localized mCherry (*nls-mCherry*), to allow identification of cells expressing Netrin (schematized in **Fig. 3A**; *ptch2:ntn1a-2A-nlsmCherry* or *ptch2:ntn1b-2A-nlsmCherry*). As a control, we injected a construct using the same promoter driving only *nls-mCherry* (*ptch2:nls-mCherry*). Each DNA construct was injected into wild-type embryos with *Tol2* transposase RNA to catalyze genomic insertion.

**Figure 3.**
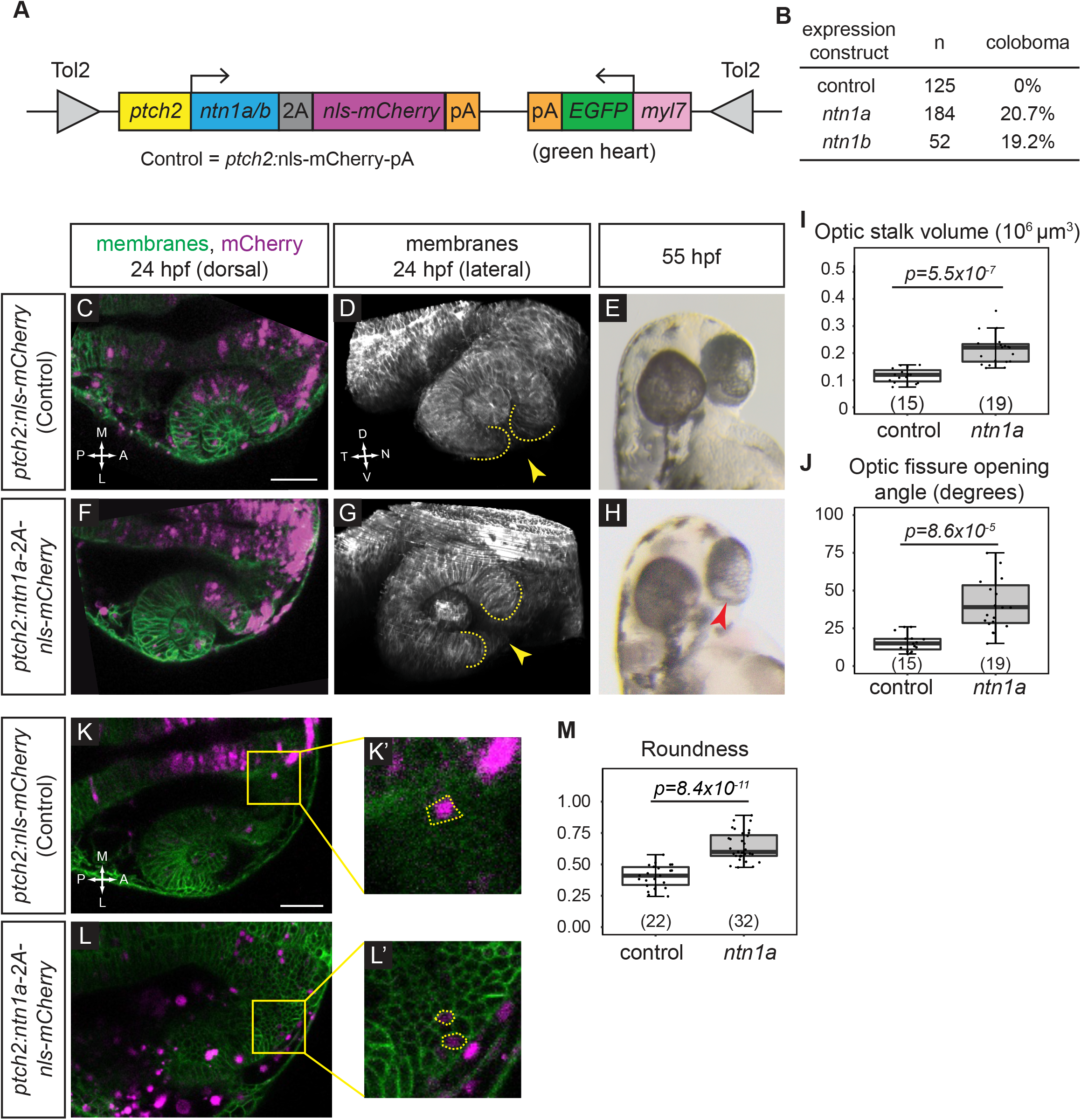
*netrin* overexpression is sufficient to disrupt cause coloboma, disrupt optic fissure formation, and perturb optic stalk cell morphology. (A) Schematic illustrating transient transgenesis expression construct (*GBS-ptch2*:*ntn1a(or ntn1b)*-2A-nls-mCherry). The control construct drives expression of only nls-mCherry. (B) Quantification of coloboma in wild-type embryos injected with the control construct, the *ntn1a*, or *ntn1b* overexpression construct. Transient transgenic overexpression of *ntn1a* or *ntn1b* is sufficient to cause coloboma in 20.7% or 19.2% of wild-type embryos, respectively. (C-M) Analysis of optic fissure formation and optic stalk cell morphology. (C-E) Wild-type embryo injected with control (nls-mCherry) transgene expression construct. (F-H) Wild-type embryo injected with the *ntn1a* overexpression transgene expression construct. (C, F) Single optical sections, dorsal view, 24 hpf. Green, membranes (Tg(*bactin2:EGFPCAAX*)); magenta, nuclei (nls-mCherry from the transgene expression construct). (D, G) Lateral views of 3D renderings show optic fissure margins (dotted lines) and opening (yellow arrowhead), 24 hpf. Grayscale, membranes only. (E, H) Optic fissure phenotypes at 55 hpf. The control embryo optic fissure is largely fused (E); (H) shows a representative *ntn1a*-overexpressing embryo with coloboma (red arrowhead indicates open, unfused fissure). (I-J) Quantification of optic stalk volume (I) and optic fissure opening angle (J), both of which are significantly increased in *ntn1a*-overexpressing embryos. (K-M) Analysis of optic stalk cell morphology, 24 hpf, dorsal view, single optical section. (K, K’) Wild-type embryo injected with control (nls-mCherry) transgene expression construct. (L, L’) Wild-type embryo injected with experimental (*ntn1a* overexpression) transgene expression construct. Green, membranes (Tg(*bactin2:EGFPCAAX*)); magenta, nls-mCherry from the transgene expression construct. (K’, L’) Zoomed views of individual transgene-expressing cells in the optic stalk, as marked by nls-mCherry fluorescence. Dotted lines show cell morphology, as visualized with (Tg(*bactin2:EGFPCAAX*)). (M) Quantification of cell elongation using the Roundness metric; *netrin1a*-overexpressing optic stalk cells are significantly less elongated than their control counterparts. Numbers in parentheses at base of graphs indicate *n*. Scale bar, 50 µm.

We first examined embryos for coloboma at 55 hpf, when the eye is pigmented and the optic fissure is nearly fused in wild-type embryos. When we assay coloboma following injection of the control construct, *ptch2:mCherry*, we observe no instances of coloboma in wild-type embryos (0/125). When *ntn1a* is overexpressed in Hh-responsive cells using the *ptch2:ntn1a* DNA construct, we observe coloboma in 20.7% (38/184) of injected embryos; injection of the *ptch2:ntn1b* DNA construct results in 19.2% (10/52) of injected embryos with coloboma (**Fig. 3B**). Although lower than the *ptch2* mutant (60-100% coloboma), this penetrance is nonetheless striking: since this experimental strategy utilizes transient overexpression, the number of transgenic cells and level of overexpression can vary widely between individual embryos. This suggests that even mosaic overexpression of *ntn1a* or *ntn1b* can cause coloboma. Since the phenotypes caused by *ntn1a* or *ntn1b* overexpression appeared similar, for the sake of simplicity, we continued our experiments using the *ptch2:ntn1a* DNA construct only.

Despite the appearance of coloboma, such a phenotype could be caused by disruption of many processes linked to optic fissure development. To examine the phenotype more closely, we asked whether *netrin* overexpression can disrupt optic fissure and stalk formation, similar to the *ptch2* mutant. Therefore, we assayed embryos at 24 hpf for optic fissure and stalk formation. Using a transgenic line to label cell membranes and provide tissue morphology, *Tg(bactin2:EGFP-CAAX)^z200^*^11^, we imaged the optic cup at 24 hpf in live embryos injected with the control (*ptch2:nls-mCherry*) or experimental (*ptch2:ntn1a-2A-nls-mCherry*) constructs (**Fig. 3C, D, F, G**). In both control and experimental embryos, nls-mCherry is largely restricted to Hh-responding cells in the brain and eye ^11^, although ectopic expression is also seen, as expected in transient transgenic experiments (**Fig. 3C, F**). Lateral views of three-dimensional renderings reveal optic fissure morphology: at the ventral side of the control (*ptch2:nls-mCherry*) eye, the optic fissure is visible as a narrow cleft with closely apposed tissue margins (**Fig. 3D**, yellow arrowhead), whereas the *ntn1a*-overexpressing optic fissure appears wide and open (**Fig. 2G**, yellow arrowhead). We quantified optic stalk and optic fissure formation, as, at optic cup stage, overactive Hh signaling results in a larger optic stalk volume and wider optic fissure opening angle than wild-type ^11^. Optic stalk volume is significantly increased in embryos overexpressing *ntn1a* compared to control embryos overexpressing mCherry (**Fig. 3I**; control 0.117 ± 0.007×10^6^ µm^3^; ntn1a overexpression 0.213 ± 0.013×10^6^ µm^3^), and optic fissure opening angle is significantly larger in embryos overexpressing *ntn1a* compared to control embryos (**Fig. 3J**; control 15.53 ± 1.585°; *ntn1a* overexpression 43.74 ± 5.590°). Both of these phenotypes are reminiscent of the *ptch2* mutant ^11^ (optic stalk volume 0.51 ± 0.08×10^6^ µm^3^; and optic fissure opening angle 59.6 ± 6.2°), although both phenotypes are quantitatively less severe in this *ntn1a*-overexpression condition, as might be expected for transient transgenesis. To determine whether embryos with increased optic stalk volume or optic fissure opening angle go on to exhibit coloboma, we raised the imaged embryos to screen for coloboma at 55 hpf. Embryos injected with the *ntn1a* construct that display a 24 hpf optic stalk and fissure phenotype go on to exhibit coloboma by 55 hpf with incomplete penetrance, again similar to the *ptch2* mutant **(Fig. 3E, H)**, a phenotype that is not observed in the control injections.

Taken together, these data suggest that spatiotemporally regulated overexpression of *netrin* is sufficient to cause coloboma, and the phenotype is similar to overactive Hh signaling in the *ptch2* mutant, with disrupted optic stalk and fissure formation.

### netrin1a overexpression disrupts single cell morphology in the optic stalk

We took these observations of aberrant tissue morphology a step further: we previously found that in the *ptch2* mutant, cell morphology in the optic stalk is disrupted ^11^. Optic stalk cells typically exhibit an elongated morphology, while in the *ptch2* mutant, optic stalk cells are significantly rounder and less elongated. We therefore asked whether the morphology of *ntn1a*-overexpressing optic stalk cells is affected similarly. We quantified optic stalk cell elongation using the metric “roundness”, a measure of aspect ratio, in Fiji ^39^, in our transient transgenic embryos (**Fig. 3K-L’**, yellow dashed outline indicates cell’s perimeter). We find that compared to control, mCherry-positive optic stalk cells are less elongated when overexpressing *ntn1a* (**Fig. 3M**; 0 represents infinitely elongated and 1 represents a perfect circle; control 0.40 ± 0.09; *ntn1a* overexpression 0.64 ± 0.11). Our observations suggest that overexpressing *ntn1a* in a subset of cells in wild-type embryos is sufficient to disrupt the optic stalk in a manner similar to loss of *ptch2*.

### Netrin ligands are not required for the ptch2 mutant coloboma phenotype

Having established that overexpression of *netrin* in wild-type embryos is sufficient to reproduce the *ptch2* mutant phenotype, we sought to determine whether Netrin is necessary for the coloboma phenotype in *ptch2* mutants. We tested if Netrin is required for the *ptch2* mutant phenotype using a genetic epistasis approach: we acquired a stable loss-of-function mutant allele for *ntn1b* ^40^, and we additionally targeted *ntn1a*, *ntn2,* and *ntn5* using the Alt-R CRISPR-Cas-9 system, which enables efficient editing in F0 injected embryos ^41,42^. Because we did not find *ntn4* upregulated in *ptch2*^-/-^ embryos, we did not test the requirement for this gene. All injected embryos were quantitatively analyzed for mutagenesis using capillary gel electrophoresis ^43^; only embryos with >70% alleles mutated for *ntn1a*, *ntn2*, and *ntn5* were included in phenotypic scoring.

To evaluate effects on eye development, we scored embryos for coloboma at 55 hpf (**Fig. 4A-E**). As expected, loss of *ptch2* results in coloboma with incomplete penetrance (**Fig. 4B, E**; 91.67±8.33%). Removal of *ntn1b* in the *ptch2* mutant background resulted in coloboma, with no change in penetrance of the phenotype (**Fig. 4C, E**; 91.67±8.33%). Finally, to remove the complete suite of upregulated Netrins (**Fig. 2G**), we injected guides against *ntn1a*, *ntn2*, and *ntn5* in the *ptch2; ntn1b* compound mutant background. We observed a similar penetrance of coloboma among uninjected and injected *ptch2; ntn1b* mutants, despite mutation of *ntn1a*, *ntn2*, and *ntn5* (**Fig. 4D, E**; 78.57±21.43%). This result suggests that Netrin is not required for the coloboma phenotype resulting from overactive Hh signaling via the *ptch2* mutant. Because these are CRISPR-injected embryos with >70% alleles mutated for all three genes, there is the possibility that some unmutated cells could provide sufficient residual Netrin activity. Additionally, it has been reported that loss of Netrins can itself result in coloboma at 2 dpf ^27,28^; this could confound a potential rescue of the *ptch2* mutant phenotype at earlier (optic cup) stages. Therefore, we sought to determine whether loss of Netrin might rescue *ptch2* mutant optic stalk and fissure phenotypes at optic cup stage (24 hpf).

**Figure 4.**
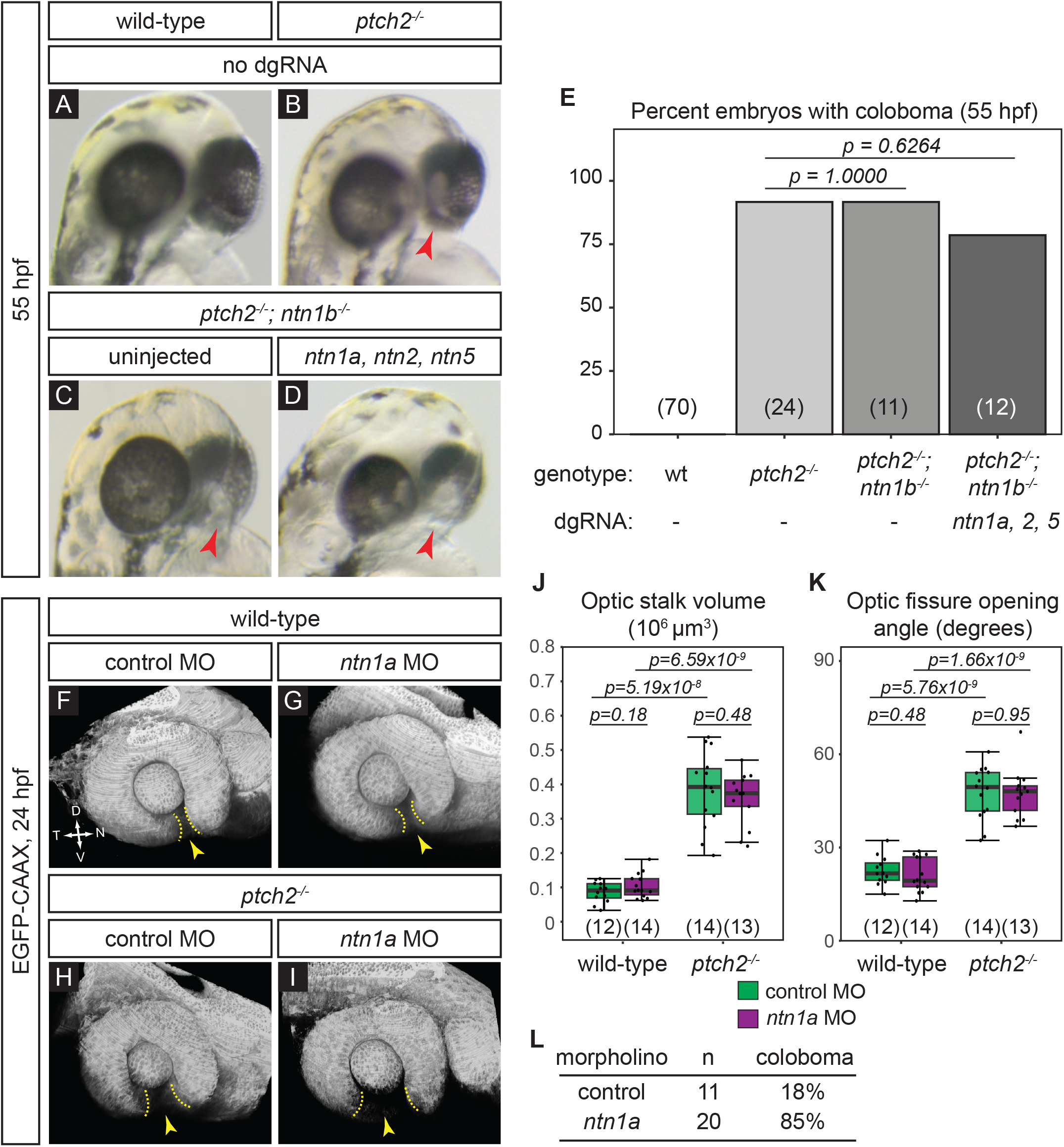
Netrin ligands are not required for the *ptch2^-/-^* coloboma phenotype. (A-E) Loss of *netrin* genes using CRISPR mutagenesis does not rescue the *ptch2^-/-^*coloboma phenotype. All embryos were evaluated for coloboma, genotyped, and gRNA-injected embryos were then individually quantified for mutagenesis efficiency. For gRNA-injected embryos, only *ptch2 ^-/-^*;*ntn1b^-/-^* embryos with >70% alleles mutated for the remaining 3 genes were analyzed. (A) wild-type (wt), (B) *ptch2^-/-^*, (C) *ptch2^-/-^*; *ntn1b^-/-^*, (D) *ptch2^-/-^*; *ntn1b^-/-^* injected with *ntn1a*, *ntn2*, and *ntn5* dgRNA, 55 hpf. Red arrowheads, coloboma. (E) Percentage of embryos with coloboma. Numbers in parentheses at base of graph indicate *n*. (F-L) Morpholino-mediated knockdown of *ntn1a* does not rescue the *ptch2^-/-^*optic fissure and stalk phenotypes. (F) wild-type injected with control MO; (G) wild-type injected with *ntn1a* MO; (H) *ptch2^-/-^* injected with control MO; (I) *ptch2^-/-^*injected with *ntn1a* MO. Tissue morphology visualized with *Tg(bactin2:EGFP-CAAX)* transgene (grayscale), 24 hpf. Yellow dotted lines, optic fissure margins; yellow arrowheads, optic fissure opening. (J, K) Quantification of optic stalk volume (J) and optic fissure opening angle (K) at 24 hpf, neither of which is affected by *ntn1a* MO injection. (L) Quantification of coloboma in embryos from *ptch2*^+/-^; *Tg(bactin2:EGFP-CAAX)* incross injected with the control MO, or the *ntn1a* MO. Knockdown of *ntn1a* in the *ptch2*^-/-^ results in a coloboma phenotype in 85% of embryos. n = total number of embryos screened.

We sought a second alternative method to impair netrin, as a complement to the CRISPR strategy, therefore, we turned to a morpholino oligonucleotide (MO) knockdown approach. We knocked down the *netrin* gene expressed in the optic fissure, *ntn1a* (**Fig. 1, 2**), using a previously validated translation-blocking *ntn1a* morpholino ^27,28^. ntn1a MO or standard control MO (0.5 pmol) was injected into embryos from a *ptch2* heterozygous incross carrying the *Tg(bactin2:EGFP-CAAX)^z200^* transgene to label cell membranes, allowing evaluation of eye morphology. Morpholino oligonucleotide efficacy was evaluated by scoring some embryos for coloboma at 52-55 hpf: as expected from prior reports ^27,28^, 17/20 (85%) *ntn1a* MO-injected embryos displayed coloboma, whereas 2/11 (18%) control MO-injected embryos displayed coloboma, within the range expected for a *ptch2* heterozygous incross **(Fig. 4L)**. Embryos were imaged at 24 hpf: optic stalk volume and optic fissure opening angle were measured, and embryos subsequently genotyped for *ptch2*. We find that in wild-type embryos, control or ntn1a MO injection has no effect on optic stalk volume (**Fig. 4F, G, J**; control MO 0.09 ± 0.03×10^6^ µm^3^; ntn1a MO 0.1 ± 0.036×10^6^ µm^3^). Optic fissure opening angle is also unaffected (**Fig. 4F, G, K**; control MO 22.5 ± 4.7°; ntn1a MO 21.1 ± 5.5°). *ptch2* mutant embryos have a large optic stalk volume (**Fig. 4H, J**; control MO 0.385 ± 0.11×10^6^ µm^3^); this is unaffected by injection of ntn1a MO (**Fig. 4I, J**; ntn1a MO 0.359 ± 0.078×10^6^ µm^3^). Similarly, injection of ntn1a MO did not affect the larger optic fissure opening angle (**Fig. 4H, I, K**; *ptch2^-/-^*+control MO 47.62 ± 8.5°; *ptch2^-/-^*+ntn1a MO 47.42 ± 7.86°).

Taken together, these data suggest that reduced *netrin* expression, either via genome editing or morpholino knockdown, does not rescue the *ptch2* mutant phenotype. Therefore, the most parsimonious interpretation of these data is that Netrin, while sufficient, is not solely required for the *ptch2* mutant eye phenotype.

## Discussion

Based on our prior work ^11^, our model is that overactive Hh signaling in the *ptch2* mutant acts through both cell- and non-cell-autonomous mechanisms to cause coloboma. We interpret this to mean that a combination of cell-intrinsic and intercellular signaling factors are responsible for disrupting cell movements to give rise to coloboma. While we demonstrate that *ntn1a* overexpression is sufficient to disrupt optic fissure and stalk formation, the factors required to act together to produce the *ptch2* mutant coloboma phenotype are still unknown.

Our model that Netrins may be sufficient but not necessary for the *ptch2* mutant phenotype is not entirely unexpected. In the context of axon guidance, these molecules are often observed to be sufficient but not necessary ^40,44,45^. Functional redundancy and compensatory mechanisms between signaling pathways and gene regulatory networks in *in vivo* systems ensure phenotypic robustness of crucial biological processes. Our results highlight the complexity and robustness of optic fissure morphogenesis, a process that likely has similar mechanisms in place to prevent perturbations.

Understanding how Netrin genes function in normal development within these tissues, and if canonical receptors, such as DCC, Unc5, or Neogenin, are involved remains an open question. Zebrafish contain numerous Netrin receptors ^46–49^, yet there is also evidence that Netrins directly bind Integrins and Laminin to modulate cell adhesion ^12,50^. These will be important pathways to examine in the future.

In this work, we focused on Netrins as one candidate downstream target of Hh signaling in eye development, but our working model implicates additional effectors that are altered in the context of overactive Hh signaling to disrupt optic fissure formation. The strategies described in this study can be used to evaluate many other potential Hh downstream targets, and in turn, uncover new molecular mechanisms controlling optic fissure morphogenesis and impacting coloboma.

## Experimental Procedures

### Zebrafish husbandry and mutant/transgenic lines

All zebrafish (*Danio rerio*) husbandry was performed under standard conditions in accordance with University of Utah Institutional Animal Care and Use Committee (IACUC) Protocol approval (Protocol #1647). Embryos (Tu or TL strains) were raised at 28.5-30 °C and staged according to time post fertilization and morphology ^51^. Mutant lines used include: *ptch2/blowout^tc294z^* ^10,11,52,53^; *ntn1b^p210^* ^40^. Transgenic alleles used were: *Tg(bactin2:EGFP-CAAX)z200* ^11^.

For genotyping, genomic DNA was extracted from single embryos or adult fins, incubated at 95 °C in 0.05 M NaOH for 30 m, then neutralized with 1 M Tris pH 8.0. The *ptch2* locus was genotyped using an HRMA protocol ^11,54^ with the following primers: ptch2HRMA_F: 5′- CTGCACCTTCCTGGTGTGTG-3′, ptch2HRMA_R: 5′- GGTAGAAATGGATTAGAGTGAGAGGAA-3′. The *ntn1b* locus was genotyped using a CAPS assay ^55^ with the following primers: ntn1b_F: 5’- ATGATAAGGATTTTGGTAACGTGCG-3’, ntn1b_R: 5’- CTTCCCGAAAGCGGAGTTCAC-3’. Full length PCR product was run on a 3% gel to distinguish bands.

### RNA synthesis and nucleic acid injections

pCS2 template (pCS2-Transposase) was linearized with NotI-HF (R3189L, New England Biolabs) and capped RNA was synthesized using the mMessage mMachine SP6 kit (AM1340, Invitrogen). RNA was purified using the RNeasy Mini Kit (74104, Qiagen) and ethanol precipitated. For Tol2 injections, 25 pg Transposase RNA and 25 pg assembled DNA construct were co-injected into one-cell embryos. For CRISPR-Cas9 injections, 5 µM dgRNA and 5 µM Cas9 protein (1074181, Integrated DNA Technologies) were co-injected into one-cell stage embryos.

### Morpholino oligonucleotide injection

Ntn1a translation blocking morpholino (5’ - CATCAGAGACTCTCAACATCCTCGC - 3’) was used, as in ^27,28^. The standard GeneTools control morpholino was used as our control. For our experiments, 0.5 pmol (4.17 ng) was injected into one-cell stage embryos.

### In situ hybridization

Embryos were fixed at 24 hpf in 4% paraformaldehyde overnight at 4 °C and dehydrated in 100% methanol. Colorimetric *in situ* hybridization was performed as described previously ^56^. *In situ* riboprobes were synthesized from linearized templates: pBluescript II SK- (*netrin1a*), pBluescript II KS+ (*netrin1b*), pGEM-Teasy (*netrin2* and *netrin4*; ^18,19^). *Netrin5* was synthesized from a PCR fragment amplified from cDNA using primers described previously with T7 RNA polymerase ^16^.

### HCR RNA-FISH

HCR was performed using an adapted version of the publicly available protocol, “HCR RNA-FISH protocol for whole-mount zebrafish embryos and larvae” (Molecular Instruments; ^35^). For each netrin gene, one HCR RNA-FISH bundle per target RNA was ordered and contained the HCR split-initiator probe set and the HCR amplifier, and the reaction was performed with HCR RNA-FISH buffers. For all target mRNAs, amplifier B1-647 was used.

### Reverse transcription-quantitative PCR

Embryos were pooled at 24 hpf (*n*=30) and immediately homogenized using the QIAshredder (79654, Qiagen). Total RNA was then extracted using the RNeasy Mini Kit (74104, Qiagen) and stored at −80°C until use. cDNA was synthesized using the iScript cDNA Synthesis kit (1708890, Bio-Rad) following the manufacturer’s recommendations, such that 1 µg of RNA was loaded into each reaction. Three biological replicates were collected for each condition.

RT-qPCR primers were designed to span exon-exon junctions and produce amplicons of ∼100 bp in length. Primer sequences used are: netrin1a_F1: 5’- GCTGTGTTTCAGCACAGGAG-3’, netrin1a_R1: 5’- CCTTTGAGCACACAGCAGAG-3’, netrin1b_F1: 5’- GGCAAGATGAAGGTCACCA-3’, netrin1b_R1: 5’- CACCGATATGATGTTGATGG-3’, netrin2_F1: 5’- AAGAGGCCAACGAGTGCTTA-3’, netrin2_R1: 5’- CACACTCCTCCACTCTTTCG-3’, netrin4_F1: 5’- TGAGCACTATGGAGCTGACG-3’, netrin4_R1: ATTTCCCATGCGTGGATTAC-3’, netrin5_F1: 5’- GCTCCGCCTGAATATCTGTC-3’, netrin5_R1: 5’- GGAGAGGGTCTGGAAAGGAG-3’, eef1a1l1_F: 5’- CCTCTTTCTGTTACCTGGCAAA-3’, eef1a1l1_R: 5’- CTTTTCCTTTCCCATGATTGA-3’. During optimization, products were both gel analyzed and sequenced to ensure product specificity. All reactions utilized the PowerUp SYBR Green Master Mix (A25741, Applied BioSystems) and were performed on an Applied BioSystems 7900HT instrument. Cycling parameters were: 50°C (2 min) followed by 40 cycles of 95°C (2 min), 58°C (15 s), 72°C (1 min), then followed by a dissociation curve. Applied BioSystems software SDSv2.4 was used to determine cycle threshold (C_t_) values and melting curves. All reactions were performed in triplicate with a ‘no-template’ control.

RT-qPCR analysis was performed in Microsoft Excel using the ΔΔCt method ^36,57^. The relative quantity (RQ) of each gene was normalized to the reference gene *eef1a1l1* ^58^, and the normalized relative quantity (NRQ) was determined by normalizing 24 hpf *ptch2^−/−^* expression to normalized 24 hpf wild-type expression.

### Generation of transient transgenesis expression constructs

For the *GBS-ptch2:nls-mCherry* control construct, the *GBS-ptch2* promoter (p5E-GBSptch2; ^11,38^) was recombined with a middle entry clone of nuclear localized mCherry (pME-nlsmCherry) and a 3’ clone of the SV40 late polyA signal sequence (p3E-polyA) into the Tol2 transposon-flanked destination vector, pDestTol2CG2 ^37^.

For the *GBS-ptch2:ntn1a-2A-nls-mCherry* and *GBS-ptch2:ntn1b-2A-nls-mCherry* experimental constructs, the *GBS-ptch2* promoter (p5E-GBSptch2) was recombined with a middle entry clone of the open reading frames of *netrin1a* (pME-*ntn1a*) or *netrin1b* (pME-*ntn1b*) lacking a stop codon, and a 3’ clone of the PTV-2A peptide, nuclear localized mCherry and the SV40 late polyA signal sequence (p3E-2A-NLSmCherry-pA) into the *Tol2* transposon-flanked destination vector, pDestTol2CG2 ^37^.

### Coloboma scoring

Embryos were individually screened and scored for coloboma at 52-55 hpf using an Olympus SZX16 stereomicroscope. The phenotype was scored by viewing the back of the eye and focusing at the depth of the RPE; embryos that were scored as positive for coloboma had eyes that displayed an expanded region lacking pigmentation in the area of the optic nerve head either unilaterally or bilaterally. This area was distinctly wider and more open than the rest of the optic fissure that was undergoing fusion at the ventral side of the optic cup. All genetic experiments were scored blindly. Embryos were subsequently genotyped as described above.

### CRISPR-Cas9 mutagenesis

gRNA target sites were identified using the web programs CHOpCHOP (http://chopchop.cbu.uib.no). Genomic DNA sequences from Ensembl GRCz11 (http://useast.ensembl.org/Danio_rerio/Info/Index) were used for target site searches. Mutagenesis was performed using the Alt-R CRISPR-Cas9 system (Integrated DNA Technologies). CRISPR RNAs (crRNAs) were designed to target exon 1 of ntn1a (5′- CAUCCCCGUCUUCGUAAACGCGG-3′), exon 1 of ntn2 (5′-CCAACCGCAUAAUAGUACGUCGG-3′), and exon 1 of ntn5 (5′- UGGACUUUGAUAGUUCCCCUAGG-3′). The crRNA and trans-activating crRNA (tracrRNA) were annealed into a dual guide dgRNA complex at a 1:1 ratio and stored at −20°C until use. On the day of injections, the ribonucleoprotein was assembled by incubating the dgRNA complexes (25 µM of total dgRNA) with the Cas9 Nuclease 3NLS (25 µM) (1074181, Integrated DNA Technologies) at 37 °C for 5 min ^42^. An injection cocktail of 5 µM dgRNA and 5 µM Cas9 protein and was injected into one-cell stage embryos.

### Fragment analysis for capillary electrophoresis

To quantify the activity and efficiency of individual crRNAs, fragment analysis by capillary electrophoresis was used. Primers were designed to amplify an ∼80 bp region surrounding the crRNA target site for *ntn1a*, *ntn2*, and *ntn5*. The forward primer was labeled with a 5’ 6- carboxyfluorescein tag (6-FAM, Integrated DNA Technologies). The following primers were used: Ntn1a_3 crRNA 56-FAM F: 5’-CGGATCCGTGTTACGACGAGAA-3’ (+6-FAM modification), Ntn1a_3 crRNA HRMA R: 5’-CTGGACGCGCGTACTTCTTTC-3’, Ntn2 crRNA 56- FAM F: 5’-AAGTGAAGACGCTCTCGGTG-3’ (+ 6-FAM), Ntn2 crRNA HRMA R: 5’- TCTCCCGGACACACCTAGAG-3’, Ntn5 crRNA 56-FAM F: 5’-GCTTCAGAGGGCTCCAGTG-3’ (+ 6-FAM), Ntn5 crRNA HRMA R: 5’-GGACTCAACCGAATCCACCT-3’. Genomic DNA was isolated from *ptch2*^-/-^; *ntn1b*^-/-^ embryos either uninjected, or injected with *ntn1a*, *ntn2*, and *ntn5* dgRNAs. Following PCR amplification, the fragments were diluted 10-fold with distilled water. Two microliters were further processed by the University of Utah DNA Sequencing Core Facility. Capillary electrophoresis was performed on an Applied BioSystems 3730 DNA analyzer (Applied Biosystems). Collected data were analyzed with GeneMapper Software (Applied Biosystems).

Analysis was adapted from previously published methods ^43^. Fragments <70 bp in size and peaks <150 in height were removed from the analysis. Six control uninjected *ptch2*^-/-^; *ntn1b*^-/-^ samples were analyzed for each mutagenesis experiment, and the highest peak was identified as wild-type (if there was a second peak similarly substantially high, that was included), and a range of fragment sizes encompassing that peak +/- 0.5 bp made up the wild-type fragment range. For each control sample, the total height of all fragments and the total height of wild-type fragments were each determined and a ratio of wild-type/total was calculated. The average ratio was then acquired across all control samples. For injected *ptch2*^-/-^; *ntn1b*^-/-^ samples, the same steps were followed. The total height of all fragments and the total height of wild-type fragments (defined using the control wild-type range) were determined, and a wild-type ratio was calculated. Each sample’s wild-type ratio was normalized using the wild-type ratio from the control samples (injected wild-type ratio/uninjected wild-type ratio). To determine the percent of transcripts in each sample that are edited (i.e. the mutant allele frequency), the equation: 1 - normalized ratio * 100 was used. Only injected samples with mutant allele frequencies for all targeted genes >70% were used in this study.

### Imaging

For confocal imaging, both live and fixed, embryos were embedded in 1.6% low melting point agarose in E3 or PBS in PELCO glass bottom dishes (14027, Ted Pella). Images were acquired using either a Zeiss LSM710 or LSM880 laser-scanning confocal microscope. All imaging was performed with a 40x water immersion objective (1.2 NA). Datasets were acquired with the following parameters: 512×512; voxel size 0.69 × 0.69 × 2.1 µm^3^. All imaging and analyses were performed blinded to the genotype of each sample.

### Image analysis: HCR quantification

*<u>ntn1a</u>* <u>domain quantification</u>: Expression domain was quantified using volume data of HCR-stained embryos. 3D data sets were oriented in FluoRender ^59^ to achieve a lateral view. This orientation was captured in FluoRender and saved as a TIFF image. The domain of *ntn1a* expression was measured in FIJI/ImageJ using the angle tool; the vertex was positioned at the center of the lens with the rays of the angle projected to encompass the extent of the *ntn1a* signal expression domain (schematized in Fig. 2J).

*<u>ntn1a</u>* <u>and *ntn1b* fluorescence intensity quantification</u>: HCR data (3D volume datasets with embryos mounted dorsally) were quantified as normalized fluorescence intensity measured in FIJI/ImageJ. For *ntn1a*, for which expression was observed in the anterior rim of the optic cup and nasal margin of the optic fissure, a maximum intensity projection was generated through the entire depth of the region. An ellipse (2019 µm^2^) was placed to encompass the anterior rim and nasal margin, and the mean fluorescence intensity measured. An ellipse of the same size (2019 µm^2^) was placed over the dorsal eye, an internal control region, and the mean fluorescence intensity again measured. Fluorescence intensity in the anterior rim and nasal margin was normalized to the dorsal eye for each embryo; the normalized values are plotted in Fig. 2K.

For *ntn1b*, quantification was carried out in a similar manner to *ntn1a*, except that the optic stalk, where signal was observed, was quantified. A maximum intensity projection through the entire depth of the optic stalk was generated. An ellipse (3031 µm^2^) was placed around the optic stalk, and the mean fluorescence intensity measured. An ellipse of the same size (3031 µm^2^) was placed over the dorsal eye, an internal control region, and the mean fluorescence intensity again measured. Fluorescence intensity in the optic stalk was normalized to the dorsal eye for each embryo; the normalized values are plotted in Fig. 2K.

### Image analysis: Optic stalk volume

In FIJI/ImageJ, the segmentation editor was used to manually segment the optic stalk in 3D volume data sets of live embryos labeled for cell membranes (EGFP-CAAX). The optic stalk was outlined using the freehand selection tool, moving through the z-stack, slice by slice. Once the entire optic stalk was segmented, the stack labels were saved as a new tiff file. The total volume of the segmented region was measured in FluoRender using the Paint Brush tools. The entire volume was selected using the brush, and then the volume measured using the “Get Selection Size” function; output was in µm^3^, based on voxel size read in from the image data.

### Image analysis: Optic fissure opening angle

Three-dimensional data sets of live embryos labeled for cell membranes (*GFP-CAAX*) were oriented in FluoRender ^59^ for a lateral view. Using the lateral cutaway tool, we cut to the lens midpoint. This orientation was captured in FluoRender and saved as a TIFF image. The opening angle of the optic fissure was measured in Fiji using the angle tool; the vertex was positioned at the center of the lens with the rays of the angle projected to each optic fissure margin.

### Image analysis: Optic stalk cell roundness

Live embryos were mounted dorsally and TIFF images were captured containing a maximum intensity projection of 10 slices that contained one or more dispersed labeled optic cell in *mCherry.* An outline was drawn around the cell using the freehand tool in Fiji and roundness was manually calculated using the area and major axis measurement.

### Box and whisker plots

Box and whisker plots were generated using the ggplot2 package in R Studio. The lower and upper hinges correspond to the first and third quartiles. The upper whisker extends from the hinge to the largest value no further than 1.5 * IQR from the hinge, and the lower whisker extends from the hinge to the smallest value at most 1.5 * IQR of the hinge. Data beyond the end of the whiskers are called “outlying” points and are plotted individually. The line in the box represents the median.

### Statistics

For all quantifications, *P*-values were calculated using an unpaired Student’s t-test in which the means of the two comparison sets are considered statistically significant if *P < 0.05*. If the variance of the two comparison sets was significantly different, Welch’s correction was used. Throughout the manuscript, quantifications in the text are listed as mean ± standard error.

## Acknowledgments

We are grateful to Puneet Dang and Jonathan Raper for generously sharing zebrafish lines. We are also grateful to David Grunwald and Kazuyuki Hoshijima; Mike Klein (University of Utah Genomics Core); and Natasha O’Brown for technical support and advice. Thanks to the Centralized Zebrafish Animal Resource and Kwan lab members past and present for useful discussion. This work was supported by the University of Utah Developmental Biology Training Grant (NIH T32 HD007491 to S. Lusk and S. LaPotin) and the National Eye Institute/National Institutes of Health (F31 EY030758 to S. Lusk, R01 EY025378 to K.M.K.).

## Author Contributions

S. Lusk and K.M.K. were responsible for conceptualization, methodology, and investigation throughout the study, and visualization, writing, and editing of the manuscript. S. LaPotin and J.S.P. were also responsible for methodology and investigation.

## Ethics Statement

All procedures and experiments with zebrafish were approved under Protocol #1647 by the University of Utah Institutional Animal Care and Use Committee and conformed to the ARVO guidelines for the use of animals in vision research.

The authors declare no conflicts of interest.

